# The FluPRINT dataset, a multidimensional analysis of the influenza vaccine imprint on the immune system

**DOI:** 10.1101/564062

**Authors:** Adriana Tomic, Ivan Tomic, Cornelia L. Dekker, Holden T. Maecker, Mark M. Davis

## Abstract

Recent advances in machine learning have allowed identification of molecular and cellular factors that underly successful antibody responses to influenza vaccines. Results of these studies have revealed the high level of complexity necessary to establish influenza immunity, and many different cellular and molecular components involved. However, identified correlates of protection, as measured by antibody responses fail to account for the majority of vaccinated cases across ages, cohorts, and influenza seasons. Major challenges arise from small sample sizes and from analysis of only one aspect of the biology such by using transcriptome data. The objective of the current study is to create a unified database, entitled FluPRINT, to enable a large-scale study exploring novel cellular and molecular underpinnings of successful antibody responses to influenza vaccines. Over 3,000 parameters were considered, including serological responses to influenza strains, serum cytokines, cell subset phenotypes, and cytokine stimulations. FluPRINT, thus facilitates application of machine learning algorithms for data mining. The data are publicly available and represent a resource to uncover new markers and mechanisms that are important for influenza vaccine immunogenicity.

## Background and Summary

Influenza virus has a devastating societal impact, causing up to 650,000 deaths every year worldwide^1^. Vaccination can prevent influenza-like illnesses, and thus lower the risk of the virus outbreak. However, currently available vaccines do not always provide protection, even among otherwise-healthy people, leading to serious pandemics. The vaccine efficacy is measured as ability of a new seasonal influenza vaccine to prevent influenza-like illness compared to the placebo group, as defined by the US Food and Drug Administration (FDA) in their guideline for vaccine licensure^2^. Young children and elderly, due to high susceptibility to influenza infection^3^, are vaccinated annually and thus, placebo-controlled clinical efficacy study in this population cannot be performed. The alternative approach to correlate vaccine-mediated protection in these populations is based on immunogenicity endpoints, recommended by FDA. The appropriate immunogenicity endpoint is the influenza-specific antibody titer measured by a hemagglutination inhibition (HAI) assay to each viral strain included in the vaccine. Vaccine protection is then assessed based on seroconversion (4-fold increase in the HAI antibody titers after vaccination) and seroprotection (geometric mean HAI titer ≥40 after vaccination). The HAI titer ≥40 after vaccination is associated with a 50% reduction in risk of influenza infection or disease^4^.

Lack of pre-existing influenza immunity, especially T cells, has been identified as one of the major predispositions for failure to generate antibody response to vaccination^5–7^. However, exact phenotypes of CD4^+^ and CD8^+^ T cells, which are important for protective influenza immunity in general and to vaccination with live attenuated influenza vaccine (LAIV) in specific, remain elusive. Application of computational biology and machine learning to clinical datasets holds promise for identifying immune cell populations and genes that mediate HAI antibody responses to influenza vaccines as a correlate of vaccine protection^8–15^. Identified correlates of protection, moreover, are not consistent between cohorts and study years^8,9,11,12^. Some of the identified challenges leading to such discrepancy are small sample sizes and analysis of only one aspect of the biology, such as molecular correlates of protection by using transcriptome data^16^. Additionally, comparison of the results of different predictive models is hampered by the lack of a consensus regarding what defines the outcome of vaccination, i.e. high vs. low responders. For these reasons, it is necessary to generate a unified dataset that includes multiple measurements across age, gender and racially diverse populations, including different vaccine types. Specifically, it is of the utmost importance to include single-cell analysis at the protein level, such as mass cytometry combined with multiple high-dimensional biological measurements, since these have power to reveal heterogeneity of the immune system^17–21^.

To accomplish that goal, we generated FluPRINT, a dataset consisting of 13 data types in standardized tables on blood and serum samples taken from 740 individuals undergoing influenza vaccination with inactivated (IIV) or live attenuated seasonal influenza vaccines (LAIV) (Fig. 1). The FluPRINT dataset contains information on more than 3,000 parameters measured using mass cytometry (CyTOF), flow cytometry, phosphorylation-specific cytometry (phospho-flow), multiplex cytokine assays (multiplex ELISA), clinical lab tests (hormones and complete blood count), serological profiling (HAI assay) and virological tests. In the dataset, vaccine protection is measured using HAI assay, and following FDA guidelines individuals are marked as high or low responders depending on the HAI titers after vaccination. The FluPRINT represents fully integrated and normalized immunology measurements from eight clinical studies conducted between 2007 to 2015 at the Human Immune Monitoring Center (HIMC) of Stanford University. Among those, one contains data from 135 donors enrolled in the 8-year long ongoing longitudinal study following immune responses to seasonal inactivated influenza vaccines. This is particularly interesting set of data that can deepen our understanding how repeated vaccination effects vaccine immunogenicity. The MySQL database containing this immense dataset is publicly available online (www.fluprint.com). The dataset represents a unique source in terms of value and scale, which will broaden our understanding of immunogenicity of the current influenza vaccines.

**Figure 1.**
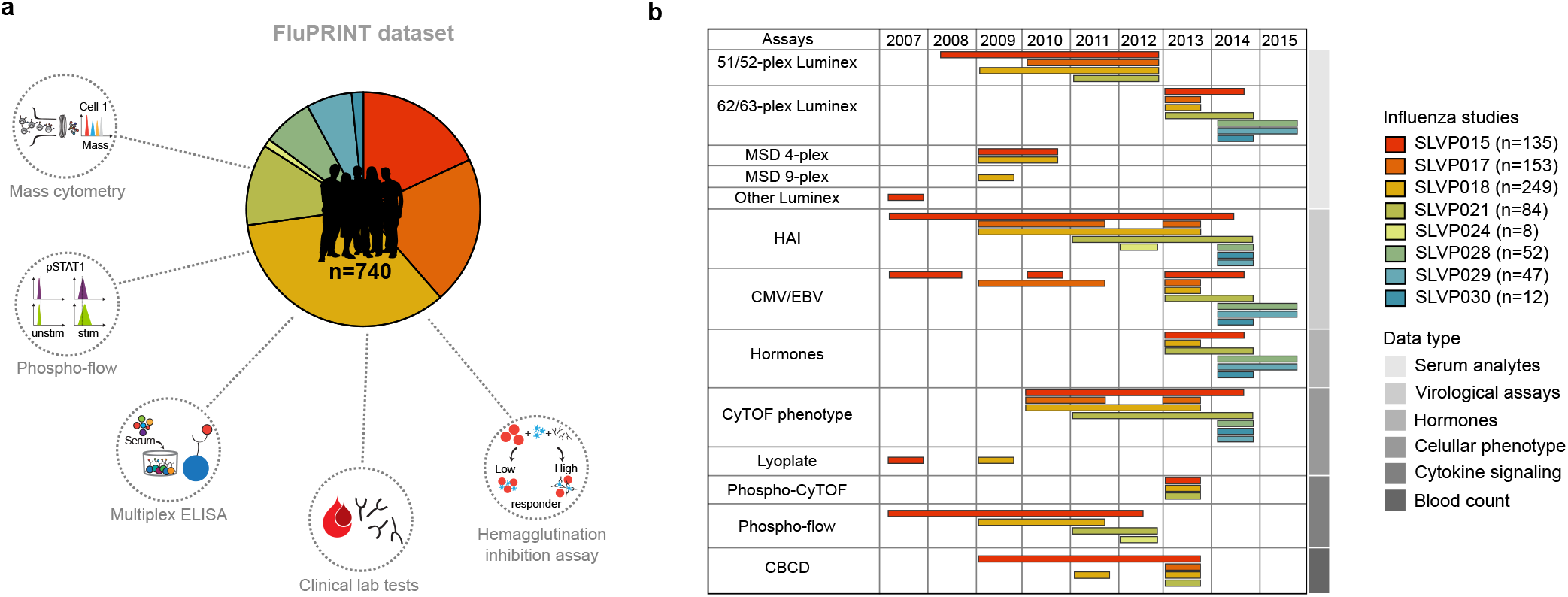
Overview of the FluPRINT dataset. The FluPRINT dataset consists of the 740 individuals from 8 clinical studies (SLVP015, SLVP017, SLVP018, SLVP021, SLVO024, SLVP028, SLVP029 and SLVP030) and 8 influenza seasons (from 2017 to 2015). (**a**) Pie chart shows distribution of donors across clinical studies. The dataset contains harmonized data from different assays, including mass and flow cytometry, phosphorylated cytometry (Phospho-flow), multiplex ELISA (Luminex assay), clinical lab tests, such as complete blood test, analysis of hormones and virological assays (CMV and EBV antibody titers) and serological profiling with hemagglutination inhibition assay, which was used to define high and low responders. (**b**) Distribution of assays across years available for each clinical study.

## Methods

### Clinical studies

All studies were approved by the Stanford Institutional Review Board and performed in accordance with guidelines on human cell research. Peripheral blood samples were obtained at the Clinical and Translational Research Unit at Stanford University after written informed consent/assent was obtained from participants. Samples were processed and cryopreserved by the Stanford HIMC BioBank according to the standard operating protocols available online at the HIMC website (https://iti.stanford.edu/himc/protocols.html).

### Data collection

Data involving individuals enrolled in influenza vaccine studies at the Stanford-LPCH Vaccine Program was accessed from the Stanford Data Miner (SDM) which holds data processed by HIMC from 2007 up to date^22^. The FluPRINT cohort was assembled by filtering the SDM for assays available in studies involving influenza vaccination. This resulted in a dataset containing data from 740 healthy donors enrolled in influenza vaccine studies at the Stanford-LPCH Vaccine Program from 2007 to 2015 in the following studies: SLVP015, SLVP017, SLVP018, SLVP021, SLVP024, SLVP028, SLVP029 and SLVP030. Table 1 provides a summary of all studies including information about clinical trial identification numbers on www.clinicaltrials.gov, clinical protocols, ImmPort accession numbers to access raw data and quality reports, and finally references to published works where data was used. ImmPort is a web portal that contains data from NIAID-funded immunology studies and clinical trials (https://immport.niaid.nih.gov/)^23^. All data contained in the FluPRINT dataset are made freely available through the Shared Data Portal on ImmPort repository. In all studies, except for study SLVP015, vaccine was administrated only once. The study SLVP015 was longitudinal study where 135 participants received vaccine in consecutive years from 2007-2015. In all studies, healthy participants were included, and in some studies (SLVP017 for the 2010, 2011 and 2013, SLVP021 and SLVP029) those that were vaccinated in the prior influenza season were excluded. A total of 121 CSV files containing processed data from various assays and studies were downloaded from SDM. The link to the 121 CSV files is provided on Zenodo^24^. Table 2 provides a summary of the demographic characteristics of the FluPRINT study population. The population spans a wide age range, from a 1-year-old to a 90-year-old, with a median age of 27 years. Among 740 individuals with available experimental data, 446 were females and 294 males. The majority (491) of the individuals were of European ancestry. The complete demographic information is available on the Zenodo^25^. Individuals were stratified into high and low responders, depending on their HAI antibody titers measured before and after vaccination, as described below. Figure 2 shows demographic information for the FluPRINT study population, including gender, ethnicity, cytomegalovirus (CMV) status, and age stratified by the outcome to vaccination. Out of 363 individuals with measured HAI responses, 111 were identified as high responders and 252 as low responders. Overall, no major differences in the gender, ethnicity distribution, or CMV status (Fig. 2a) or age (Fig. 2b) were observed between high and low responders.

**Table 1.** Characteristics of the clinical studies included in the FluPRINT dataset.

| Stanford study ID | ClinicalTrials.gov ID | Name | Description | Vaccines | Data in FluPRINT | ImmPort ID (www.immport.org) | Ref. |
| --- | --- | --- | --- | --- | --- | --- | --- |
| SLVP015 | NCT01827462 | Comparison of immune responses to influenza vaccine in adults of different ages (2007 - 2017) | **Who:** 18-100yo healthy participants<br>**How:** immunized annually with the seasonal inactivated influenza vaccines from 2007-2017<br>**When:** Blood samples acquired before immunization (Day 0), on days 6-8 and 28 after immunization | **2007-2013**<br>Seasonal trivalent, inactivated influenza vaccines (Fluzone)<br>**2014-2015**<br>High Dose trivalent Fluzone for participants $\geq 65$yo and quadrivalent Fluzone for younger participants | **135 donors**<br>Assays:<br>51-plex<br>Luminex<br>62-plex<br>Luminex<br>MSD 4plex<br>MSD9plex<br>Other<br>Luminex<br>HAI<br>CMV/EBV<br>Hormones<br>CyTOF<br>phenotype<br>Lyoplate<br>Phospho<br>Cytof pheno<br>Phospho<br>cytof<br>phospho<br>Phosphoflow<br>CBCD | SDY887 (2007)<br>SDY212 (2008)<br>SDY312 (2009)<br>SDY311 (2010)<br>SDY112 (2011)<br>SDY315 (2012)<br>SDY478 (2013)<br>SDY1464 (2014) | 8-10,28,37-46 |
| SLVP017 | NCT02133781<br>NCT03020498<br>NCT03020537 | B-cell immunity to influenza (2009- 2011 and 2013) | **Who:**<br>1-2yo (2013), 8-100yo healthy participants who did not receive the seasonal influenza | **2009-2011**<br>Seasonal trivalent, inactivated influenza vaccines (Fluzone) or seasonal live, | **153 donors**<br>Assays:<br>51-plex<br>Luminex | SDY1467(2009)<br>SDY1468(2010)<br>SDY1469(2011)<br>SDY1470(2012)<br>SDY1471(2013) | 27,47-50 |
|  |  |  | vaccine in previous years (2010, 2011 and 2013)<br>**How:** immunized with either seasonal inactivated or live, attenuated influenza vaccines in 2009, 2010, 2011 and 2013<br>**When:** Blood samples acquired before immunization (Day 0) and on day 28 after immunization | attenuated influenza vaccine (FluMist)<br>**2013**<br>Seasonal trivalent inactivated influenza vaccine- (Fluzone) - pediatric formulation for 1-2yo children | 62-plex<br>Luminex<br>HAI<br>CMV/EBV<br>CyTOF<br>phenotype<br>CBCD |  |  |
| SLVP018 | NCT01987349<br>NCT03022396<br>NCT03022422<br>NCT03022435<br>NCT03023176 | T-cell and general immune response to seasonal influenza vaccine (2009-2013) | **Who:** 1-8yo (2013), 8-100yo healthy participants<br>**How:** immunized with either seasonal inactivated or live, attenuated influenza vaccines from 2009-2013<br>**When:** Blood samples acquired before immunization (Day 0), days 7-10 and 28 after immunization | **2009-2010**<br>Seasonal trivalent inactivated influenza vaccine (Fluzone) or seasonal trivalent live attenuated influenza vaccine (FluMist)<br>**2010**<br>High Dose trivalent Fluzone for participants ≥ 65yo<br>**2013**<br>Seasonal trivalent, inactivated influenza Pediatric Dose | **249 donors**<br>Assays:<br>51-plex<br>Luminex<br>62-plex<br>Luminex<br>MSD 4plex<br>MSD 9plex<br>HAI<br>CMV/EBV<br>Hormones<br>CyTOF<br>phenotype<br>Lyoplate<br>Phospho<br>Cyt of pheno | SDY514(2009)<br>SDY515(2010)<br>SDY519(2011)<br>SDY1465(2012)<br>SDY1466(2013) | 27,44,51-57 |
|  |  |  |  | (Fluzone, 0.25ml) for 1-8yo children | Phospho<br>cytof<br>phospho<br>Phosphoflow<br>CBCD |  |  |
| SLVP021 | NCT02141581 | Plasmablast trafficking and antibody response in influenza vaccination (2011-2014) | **Who:** 8-34yo healthy participants who did not receive the seasonal influenza vaccine in previous years<br>**How:** immunized with either seasonal inactivated influenza vaccines given intramuscularly or intradermally and live, attenuated influenza vaccines from 2011-2014<br>**When:** Blood samples acquired before immunization (Day 0), days 6-8 and 24-32 after immunization | **2011-2014**<br>Seasonal trivalent inactivated influenza vaccine (Fluzone) given either intramuscularly or intradermally<br>**2011-2012**<br>Seasonal trivalent live attenuated influenza vaccine (FluMist) | **84 donors**<br>Assays:<br>51-plex<br>Luminex<br>62-plex<br>Luminex<br>HAI<br>CMV/EBV<br>Hormones<br>CyTOF<br>phenotype<br>Phospho<br>Cytof pheno<br>Phospho<br>cytof<br>phospho<br>Phosphoflow<br>CBCD | SDY113 (2011)<br>SDY305 (2012)<br>SDY472 (2013)<br>SDY1479 (2014) | 58-60 |
| SLVP024 | NCT03023683 | Protective mechanisms against a pandemic respiratory virus (2012) | **Who:** 2-49yo healthy participants<br>**How:** immunized with the seasonal live, attenuated influenza vaccine | Seasonal live, attenuated influenza vaccine (FluMist) | **Donors: 8**<br>Assays:<br>HAI<br>Phosphoflow | SDY1472 |  |
|  |  |  | **When:** Blood samples only from 18-42yo adults acquired before immunization (Day 0), days 7 and 28 after immunization |  |  |  |  |
| SLVP028 | NCT03088904 | Genetic and environmental factors in the response to influenza vaccination (2014-2018) | **Who:** 12-49yo healthy participants<br>**How:** immunized with either seasonal inactivated or live, attenuated influenza vaccines from 2014-2018<br>**When:** Blood samples acquired before immunization (Day 0), days 6-8 and 28+7 after immunization | Seasonal quadrivalent inactivated influenza vaccine (Fluzone) or seasonal quadrivalent live attenuated influenza vaccine (FluMist) | **Donors: 52**<br>Assays:<br>62-plex<br>Luminex<br>HAI<br>CMV/EBV<br>Hormones<br>CyTOF<br>phenotype | SDY1480 (2014)<br>SDY1481 (2015) |  |
| SLVP029 | NCT03028974 | Innate and acquired immunity to influenza infection and immunization (2014-2017) | **Who:** 6 mo-49yo healthy participants (who did not receive LAIV in the prior season nor received influenza immunizations in two or more prior seasons)<br>**How:** immunized with either seasonal<br>inactivated or live, attenuated influenza vaccines from 2014-2017<br>**When:** Blood samples acquired before immunization (Day 0), days 7 and 28 after immunization | Seasonal quadrivalent inactivated influenza vaccine (Fluzone) or seasonal quadrivalent live attenuated influenza vaccine (FluMist) | **Donors: 47**<br>Assays:<br>62-plex<br>Luminex<br>HAI<br>CMV/EBV<br>Hormones<br>CyTOF<br>phenotype | SDY1482 (2014)<br>SDY1483 (2015) |  |
| SLVP030 | NCT03453801 | The role of CD4+ memory phenotype, memory, and effector t cells in vaccination and infection (2014-2019) | **Who:** 6 mo-10yo healthy participants<br>**How:** immunized annually with either seasonal inactivated or live, attenuated influenza vaccines from 2014-2019<br>**When:** Blood samples acquired before immunization (Day 0), days 7 and 60 after immunization | Seasonal quadrivalent inactivated influenza vaccine (Fluzone) or seasonal quadrivalent live attenuated influenza vaccine (FluMist)<br>Seasonal trivalent, inactivated influenza Pediatric Dose (Fluzone, 0.25ml) for 6-35mo children | **Donors: 12**<br>Assays:<br>62-plex<br>Luminex<br>HAI<br>CMV/EBV<br>Hormones<br>CyTOF<br>phenotype | SDY1484 (2014) |  |

**Table 2.**
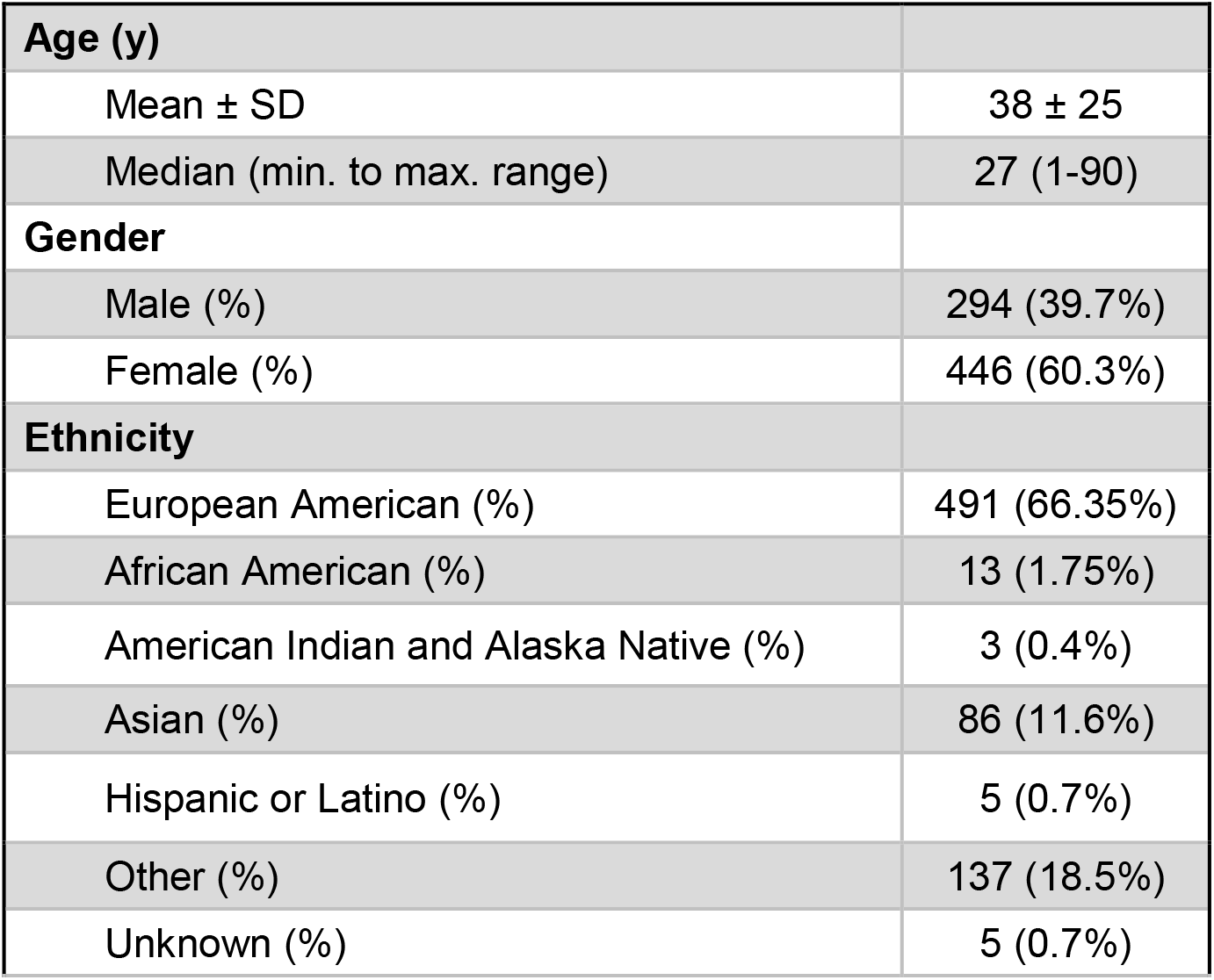
Demographic characteristics for the FluPRINT study population.

| **Age (y)** |  |
| --- | --- |
| Mean ± SD | 38 ± 25 |
| Median (min. to max. range) | 27 (1-90) |
| **Gender** |  |
| Male (%) | 294 (39.7%) |
| Female (%) | 446 (60.3%) |
| **Ethnicity** |  |
| European American (%) | 491 (66.35%) |
| African American (%) | 13 (1.75%) |
| American Indian and Alaska Native (%) | 3 (0.4%) |
| Asian (%) | 86 (11.6%) |
| Hispanic or Latino (%) | 5 (0.7%) |
| Other (%) | 137 (18.5%) |
| Unknown (%) | 5 (0.7%) |

**Figure 2.**
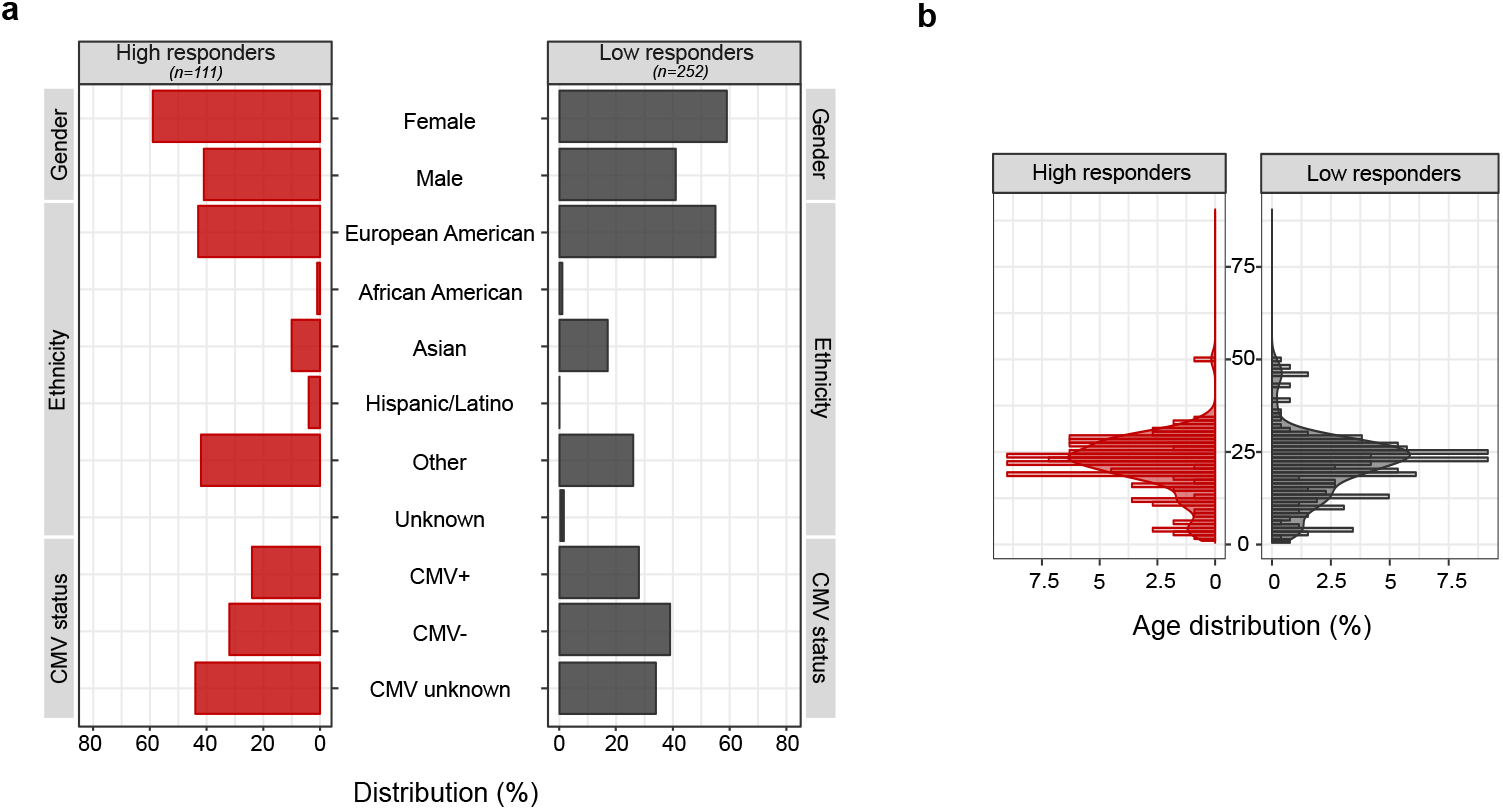
Demographic characteristics for the FluPRINT study population stratified by the vaccination outcome. Distribution of individuals in the categories of high (red, n = 111) and low (grey, n = 252) responders regarding the (**a**) gender, ethnicity and CMV status (**b**) age distribution between high and low responders. Age is indicated in years.

### Assays and data processing

All data used were analysed and processed at the HIMC^26^. The distribution of assays performed across clinical studies and years is illustrated in Fig. 1b. Overall, SLVP015 was the longest study, running from 2007 to 2014, spanning 135 unique individuals, while the majority of samples (249) came from the SLVP018 study (Fig. 1). Raw data, including report files, standards, controls, antibodies used are available at ImmPort (https://immport.niaid.nih.gov/) under identification numbers for each study provided in the Table 1. Table 3 provides information about all assays performed, protocols, validations used and references to the published manuscripts using the data. Protocols for all assays are available online at the HIMC website (https://iti.stanford.edu/himc/protocols.html).

**Table 3.** Assays performed.

| **Assays** | **Protocols** | **Validations** | **Ref.** |
| --- | --- | --- | --- |
| 51/52-plex<br>Luminex | HIMC website:<br>“Luminex-Polystyrene bead<br>kits/Luminex” | Report file available at ImmPort<br>(https://immport.niaid.nih.gov/)<br>contains all information of<br>standards, Curve fitting, Bead<br>counts to examine quality of assay<br>and CV of both samples and<br>standards. | 9,10,22,27,28 |
| 62/63-plex<br>Luminex | HIMC website<br>“Luminex –<br>eBioscience/Affymetrix<br>Magnetic bead Kits” |  |  |
| MSD 4-plex | V-PLEX Human<br>Proinflammatory Panel II<br>(Mesoscale, Cat No<br>K15053D) | Manufacturer standards | 22 |
| MSD 9-plex | Human ProInflammatory 9-Plex Ultra-Sensitive Kit<br>(Mesoscale, Cat No<br>K15007C) |  |  |
| Other<br>Luminex | Human 42-Plex<br>Polystyrene Kit<br>(EMD Millipore, H42;<br>MPXHCYTO060KPMX42) | Report file available at ImmPort<br>(https://immport.niaid.nih.gov/)<br>contains all information of<br>standards, Curve fitting, Bead<br>counts to examine quality of assay<br>and CV of both samples and<br>standards. | 28 |

| Assays | Protocols | Validations | Ref. |
| --- | --- | --- | --- |
| HAI | HIMC website<br>“Hemagglutinin inhibition (HAI) assay” | Sample, virus control, HIMC human control serum (CONS2) and control PBS available at ImmPort. | 9,10,27 |
| CMV/EBV | CMV IgG ELISA (Calbiotech, Cat No CM027G)<br>EBV-VCA IgG ELISA (Calbiotech, Cat No EVO10G) | Manufacturer standards | 10,27,30 |
| Hormones | Free Testosterone ELISA Kit (Calbiotech) and Custom Steroid Hormone Panel (human) Assay Kit (Mesosclae, MSD 4-plex) | Manufacturer standards | 8 |
| CyTOF phenotype | HIMC website<br>“CyTOF Immunophenotyping/ CyTOF phenotyping” | Data analysed using FlowJo software. Gates were adjusted on a donor-specific basis. The statistics for each gated population was exported to an Excel spreadsheet. The percentage of each cell type is determined and reported as a percent of the parent cell type. | 27,30<br>**Protocols**<br>31,32 |
| Lyoplate | HIMC website:<br>“Flow cytometry phenotyping” |  | 27,30 |
| Phospho-CyTOF (whole blood) | HIMC website<br>“Whole blood phospho-CyTOF/ Phosphoflow whole blood CYTOF” | The percentage of each cell type was determined and reported as a percent of the parent cell type. Median values were reported to quantitate the level of phosphorylation of each protein in response to stimulation. | **Protocol**<br>34 |
| Phospho-flow | HIMC website<br>“Phospho-flow-cytokine/ Phosphoepitope Flow Cytometry (Cytokine stimulation, pSTAT readouts)” |  | 9,10,27,28,30 |
| Blood count (CBCD) | Clinical haematology test performed on a Coulter counter | Performed at the Stanford Clinical Lab | - |

### Multiplex cytokine assay

Multiplex ELISA using Luminex was performed using either polystyrene bead (for 51/52-plex) or magnetic bead kits (62/63-plex) (eBioscience/Affymetrix). The processed Luminex data available in the FluPRINT is normalized at the plate level to mitigate batch and plate effects^26^. The two median fluorescence intensity (MFI) values for each sample for each analyte were averaged, and then log-base 2 transformed. Z-scores ((value–mean)/standard deviation) were computed, with means and standard deviations computed for each analyte for each plate. Thus, units of measurement were Zlog2 for serum Luminex. Part of the Luminex data was used in previous publications^9,10,22,27,28^. In 2009 and 2010, for SLVP015 and SLVP018 studies, serum analytes were analysed using MSD 4-and 9-plex kits (V-PLEX Human Proinflammatory Panel II, Mesoscale, Cat No K15053D and Human ProInflammatory 9-Plex Ultra-Sensitive Kit, Mesoscale, Cat No K15007C) as according to the manufacturer’s protocol. The assay named ‘Other Luminex’ was performed only for study SLVP015 in 2007 using the Human 42-Plex Polystyrene Kit (EMD Millipore, H42; MPXHCYTO060KPMX42) and data was processed in the same way as for the Luminex assays described above (measurement units reported were Zlog2)^28^.

### Hemagglutination inhibition assay

Serum antibody titers before vaccination and day 28 after vaccination were measured by HAI assay^29^ using strains of influenza contained in the vaccines^9,10,27^. Geometric mean titers (GMT) were calculated for all strains of the virus contained in the vaccine, while fold change is calculated as: GMT for all vaccine strains on day 28 / GMT for all vaccine strains on day 0. High responders were determined as individuals that seroconverted (4-fold or greater rise in HAI titer) and were seroprotected (GMT HAI ≥ 40).

### Virological assays

CMV and Epstein-Barr virus (EBV) analysis was performed using CMV IgG ELISA (Calbiotech, Cat No CM027G) and EBV-VCA IgG ELISA (Calbiotech, Cat No EVO10G), following manufacturer’s protocols^10,27,30^.

### Immunophenotyping

Immunophenotyping was performed either with flow cytometry (Lyoplate)^27,30^ or mass cytometry (CyTOF)^30–32^. Data was analysed using FlowJo software using the standard templates. Gates were adjusted on a donor-specific basis, if necessary, to control for any differences in background or positive staining intensity. The statistics was exported for each gated population to a spreadsheet. The percentage of each cell type is determined and reported as a percent of the parent cell type.

### Phosphorylation-specific cytometry

Phospho-flow assays were performed either using flow cytometry on PBMC (for studies SLVP015, SLVP018 and SLVP021 from 2007 to 2012)^9,10,27,28,30^ or mass cytometry on whole blood (for studies SLVP015, SLVP018 and SLVP021 in 2013)^33,34^. The percentage of each cell type is determined and reported as a percent of the parent cell type. Median values are reported to quantitate the level of phosphorylation of each protein in response to stimulation. For phospho-flow data acquired on flow cytometer a fold change value was computed as the stimulated readout divided by the unstimulated readout (e.g. 90th percentile of MFI of CD4^+^ pSTAT5 IFNα stimulated / 90th percentile of CD4^+^ pSTAT5 unstimulated cells), while for data acquired using mass cytometry a fold change was calculated by subtracting the arcsinh (intensity) between stimulated and unstimulated (arsinh stim – arcsing unstim).

### Automated importer and data harmonization

After collecting the data, a custom PHP script was generated to parse each of the 121 CSV files and to import data into the MySQL database. The source code for the script is available online at https://github.com/LogIN-/fluprint. The script optimizes the data harmonization process essential for combining data from different studies. Control and nonsense data were not imported, such as “CXCR3-FMO CD8+ T cells”, “nonNK-nonB-nonT-nonmonocyte-nonbasophils”, “viable”, etc. To standardize data, the original CSV entries were cleaned into the MySQL database readable format (e.g. quotes and parenthesis replaced with underscores, “+” with text “positive”, etc.). Additionally, classifications for ethnicity (Table 4), vaccine names (Table 5) and vaccination history (Table 6) were resolved into standard forms, while assays were numerated (Table 7). For example, “Fluzone single-dose syringe” and “Fluzone single-dose syringe 2009-2010” were mapped to “Fluzone” and given number 4 (Table 5). In all studies, vaccines were given intramuscularly for IIV and intranasally for LAIV, except for one study where IIV was given intradermally and this was labelled as Fluzone Intradermal and given number 2. During data merging, we replaced text strings with binary values. For example, for the variable of gender, female and male were replaced with zero and one. To be able to distinguish between visits in consecutive years, a unique visit identification was calculated. For the original internal visit data, each visit in one year was labelled as V1 for day zero and V2 for day seven. However, if the same individual came in the consecutive year, day zero visit would again be labelled V1, and day seven as visit V2, causing repetition of values. To avoid such repetitions in the database, we generated a unique visit ID. Therefore, for the above example, first visit in the first year would be labelled V1 for day zero and V2 for day seven, but for the next year visits would be labelled as V3 for day zero and V4 for day seven. To distinguish between Luminex assays, the prefix L50 was given to each analyte analysed with the 51/52-plex Luminex kit. Finally, we imputed new values and calculated the vaccine outcome parameter using HAI antibody titers. High responders were determined as individuals that have HAI antibody titer for all vaccine strains ≥40 after vaccination and GMT HAI fold change ≥ 4, following FDA guidelines for evaluation of vaccine efficacy^2^. Vaccine outcome was expressed as a binary value: high responders were given value of one and low responders the value zero.

**Table 4.** Remapping ethnicity.

| Original | Remapped |
| --- | --- |
| Caucasian or White | Caucasian |
| Caucasian or White,Asian | Other |
| Caucasian or White,Other | Other |
| Asian | Asian |
| Asian,Other | Other |
| Other | Other |
| Caucasian or White,Black African American,Asian,Other | Other |
| Caucasian or White,Black African American | Other |
| NULL | Other |
| Not Hispanic or Latino | Other |
| Non-Hispanic | Other |
| Decline to answer | Unknown |
| Black African American | Black or African American |
| Black African American,Asian | Other |
| Cauc or White,Black Af Am | Other |
| Caucasian or White,Pacific Islander | Other |
| Caucasian or White,Paclslan | Other |
| Cauc or White,Pacific Islander | Other |
| Pacific Islander,Asian | Other |
| American Indian/Alaska native,Caucasian or Wh | Other |
| American Indian/Alaska native,Caucasian or White | Other |
| American Indian/Alaska native,Black African American | Other |
| Am In/Alaska native,Cauc or W | Other |
| Am In/AlaskaNative,Black Af Am | Other |
| American Indian/Alaska native | American Indian or Alaska Native |
| Hispanic | Hispanic/Latino |
| Hispanic or Latino | Hispanic/Latino |

**Table 5.**
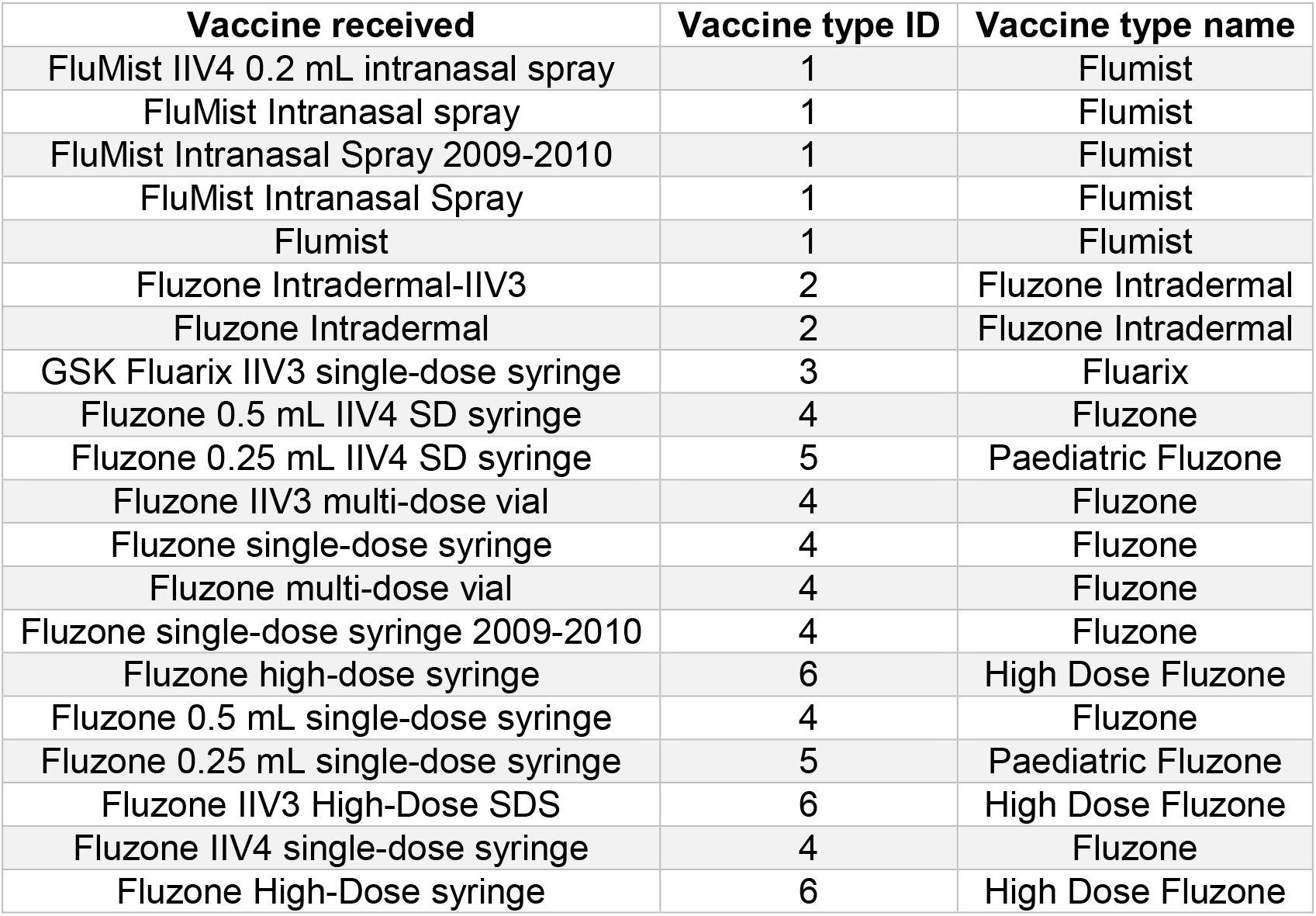
Remapping vaccine type.

| Vaccine received | Vaccine type ID | Vaccine type name |
| --- | --- | --- |
| FluMist IIV4 0.2 mL intranasal spray | 1 | Flumist |
| FluMist Intranasal spray | 1 | Flumist |
| FluMist Intranasal Spray 2009-2010 | 1 | Flumist |
| FluMist Intranasal Spray | 1 | Flumist |
| Flumist | 1 | Flumist |
| Fluzone Intradermal-IIV3 | 2 | Fluzone Intradermal |
| Fluzone Intradermal | 2 | Fluzone Intradermal |
| GSK Fluarix IIV3 single-dose syringe | 3 | Fluarix |
| Fluzone 0.5 mL IIV4 SD syringe | 4 | Fluzone |
| Fluzone 0.25 mL IIV4 SD syringe | 5 | Paediatric Fluzone |
| Fluzone IIV3 multi-dose vial | 4 | Fluzone |
| Fluzone single-dose syringe | 4 | Fluzone |
| Fluzone multi-dose vial | 4 | Fluzone |
| Fluzone single-dose syringe 2009-2010 | 4 | Fluzone |
| Fluzone high-dose syringe | 6 | High Dose Fluzone |
| Fluzone 0.5 mL single-dose syringe | 4 | Fluzone |
| Fluzone 0.25 mL single-dose syringe | 5 | Paediatric Fluzone |
| Fluzone IIV3 High-Dose SDS | 6 | High Dose Fluzone |
| Fluzone IIV4 single-dose syringe | 4 | Fluzone |
| Fluzone High-Dose syringe | 6 | High Dose Fluzone |

**Table 6.** Remapping vaccination history.

| Original | Remapped |
| --- | --- |
| No | 0 |
| Yes | 1 |
| IIV injection/im | 2 |
| Doesn't know/doesn't remember/na/does not remember | 3 |
| LAIV4 intranasal/ laiv_std_intranasal/ laiv_std_intranasal/nasal/intranasal | 4 |

**Table 7.** Assays in the database.

| Original | Remapped |
| --- | --- |
| CMV EBV | 1 |
| Other immunoassay | 2 |
| Human Luminex 62-63 plex | 3 |
| CyTOF phenotyping | 4 |
| HAI | 5 |
| Human Luminex 51 plex | 6 |
| Phospho-flow cytokine stim (PBMC) | 7 |
| pCyTOF (whole blood) pheno | 9 |
| pCyTOF (whole blood) phospho | 10 |
| CBCD | 11 |
| Human MSD 4 plex | 12 |
| Lyoplate 1 | 13 |
| Human MSD 9 plex | 14 |
| Human Luminex 50 plex | 15 |
| Other Luminex | 16 |

### Generating tables

To build FluPRINT database, we generated four tables, as shown in Figure 3. Table 8 depicts characteristics of the FluPRINT database. In the table ***donor***, each row represents an individual given a unique encrypted identification number (study donor ID). Other fields provide information about the clinical study in which an individual was enrolled (study ID and study internal ID), gender and race. The second table, named ***donor_visits*** describes information about the donor’s age, CMV and EBV status, Body Mass Index (BMI) and vaccine received on each clinical visit. Each clinical visit was given a unique identification (visit ID) in addition to the internal visit ID (provided by the clinical study) to distinguish between visits in consecutive years. For each visit, we calculated vaccine response by measuring HAI antibody response. Information about vaccine outcome is available as geometric mean titers (geo_mean), difference in the geometric mean titers before and after vaccination (delta_geo_mean), and difference for each vaccine strain (delta_single). In the last field, each individual is classified as high and low responder (vaccine_resp). On each visit, samples were analysed and information about which assays were performed (assay field) and value of the measured analytes (units and data) are stored in the ***experimental_data*** table. Finally, the ***medical_history*** table describes information connected with each clinical visit about usage of statins (statin_use) and if influenza vaccine was received in the past (influenza vaccine history), if yes, how many times (total_vaccines_received). Also, we provide information which type of influenza vaccine was received in the previous years (1 to 5 years prior enrolment in the clinical study). Lastly, information about influenza infection history and influenza-related hospitalization is provided.

**Table 8.**
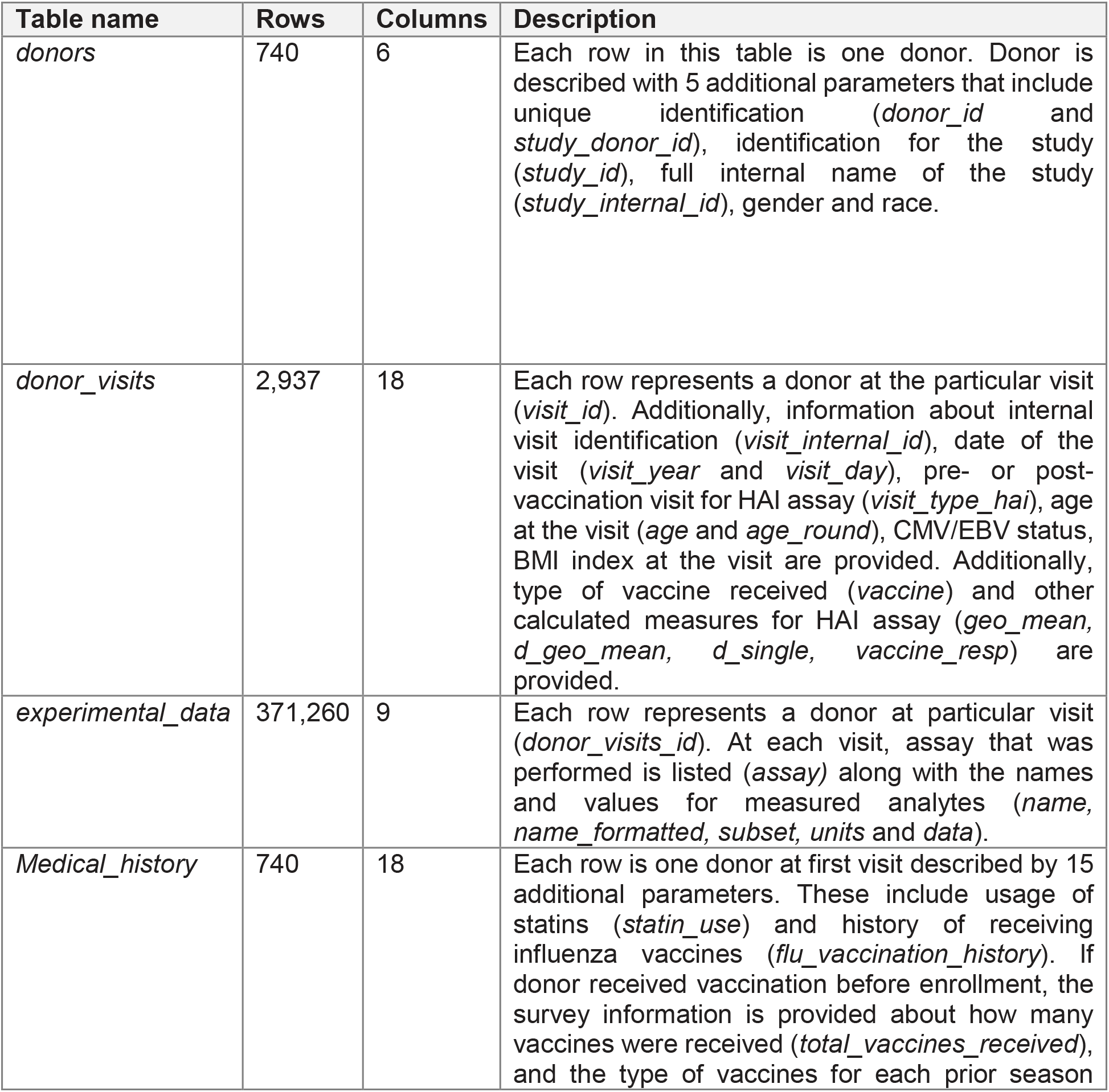
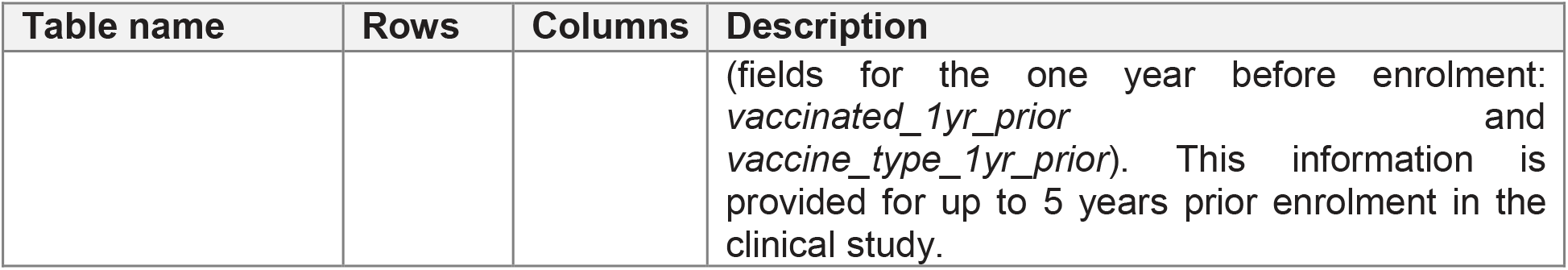
The characteristics of the FluPRINT database.

**Figure 3.**
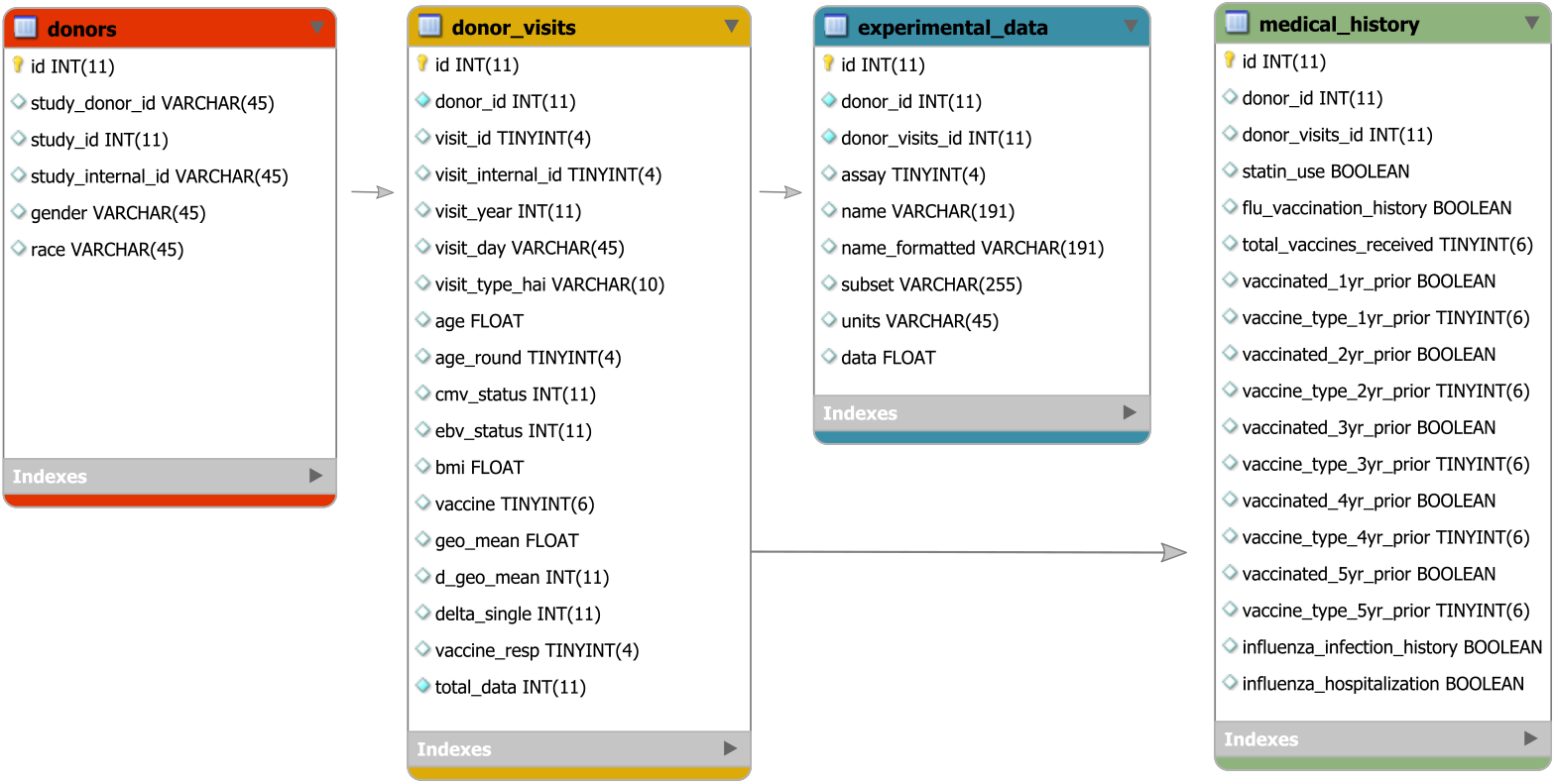
The FluPRINT database model. The diagram shows a schema of the FluPRINT database. Core tables, donors (red), donor_visits (yellow), experimental_data (blue) and medical_history (green) are interconnected. Tables experimental_data and medical_history are connected to the core table donor_visits. The data fields for each table are listed, including the name and the type of the data. CHAR and VARCHAR, string data as characters; INT, numeric data as integers; FLOAT, approximate numeric data values; DECIMAL, exact numeric data values; DATETIME, temporal data values; TINYINT, numeric data as integers (range 0-255); BOOLEAN, numeric data with Boolean values (zero/one). Maximal number of characters allowed in the data fields is denoted as number in parenthesis.

### Code availability

The source code for the PHP script and database schema are available from a public github repository (https://github.com/LogIN-/fluprint). Raw data files used to generate dataset are provided as single compressed file on Zenodo^24^. Full study population with demographic characteristics is provided as single CSV file^25^. Additionally, entire FluPRINT database export is available as CSV table and SQL file^35^. Database is also accessible at the project website https://fluprint.com.

## Data Records

The FluPRINT dataset described herein is available online for use by the research community and can be downloaded directly from a research data repository Zenodo^35^. Additionally, the dataset can be imported in the MySQL database for further manipulation and data extraction. The instructions how to import FluPRINT into the database are available at github (https://github.com/LogIN-/fluprint). The summary of the dataset, including the number of observations, fields and description for each table is provided in Table 8.

## Technical Validation

The objective of the current study was to ensure that the FluPRINT dataset accurately reflects processed data available in SDM. Technical data validation was carried in previous published studies referred in the Table 3. Data was downloaded from the original source, and here we focused on ensuring that data records were accurately harmonized, merged and mapped in the unifying FluPRINT database.

The FluPRINT dataset was validated on two levels: (1) upon insertion and (2) after the data was inserted into the database. To validate data on insertion, we created loggers to monitor import of the CSV files into the database. This ensured easier and more effective troubleshooting of potential problems and contributed to the monitoring of the import process. Two different sets were used: (1) informative and (2) error loggers. Informative loggers provided information about which processing step has started or finished and how many samples have been processed in that particular step. This allowed us to monitor that correct number of samples was processed. Error loggers provided exact identification and name of the data which could not be imported into the database, usually caused by missing or incorrect user input, such as “…assay is missing. Skipping…’$row’”. This facilitated the process to identify erroneous data, which were then manually reviewed, corrected, and updated.

Once the database was built, a manual review of data was performed to ensure accuracy and integrity of the dataset. Several random individuals were chosen and the accuracy of data was evaluated by comparison with the raw data. Additionally, we evaluated total number of all donors, assays performed, clinical studies and years with the raw data available at the SDM.

## Usage Notes

Recent advances in the computational biology and the development of novel machine learning algorithms, especially deep learning, make it possible to extract knowledge and identify patterns in an unbiased manner from large clinical datasets. Application of machine learning algorithms to clinical datasets can reveal biomarkers for different diseases, therapies^36^, including vaccinations^8,9,12^. The data from the FluPRINT study can be used to identify cellular and molecular baseline biomarkers that govern successful antibody response to influenza vaccines (IIV and LAIV) across different influenza seasons and a broad age population. The HAI antibody response to influenza vaccines is considered as an alternative way to compare efficacy of the vaccines in susceptible groups where placebo-controlled clinical efficacy study cannot be performed. Since FluPRINT dataset is provided as a database, this facilitates further analysis. Queries can be easily performed to obtain a single CSV file. For example, researchers interested in understanding which immune cells and chemokines can differentiate between high and low responders that received inactivated influenza vaccine could search the FluPRINT database. In the database, they can find all donors for which flow cytometry or mass cytometry were performed together with Luminex assays, for which donors the HAI response was measured, and all the donors who received inactivated influenza vaccine. The resulting CSV file can then easily be used for downstream analysis.

Major advantages of this dataset are the mapping of the vaccine outcome, classifying individuals as high or low responders, standardization of the data from different clinical studies, and from different assays. This data harmonization process allows for direct comparison of immune cell frequency, phenotype, and functionality and quantity of chemokines and cytokines shared between individuals before or after influenza vaccinations. By releasing the FluPRINT database and the source code, we provide users with the ability to continue building upon this resource and to update the database with their data and other databases.

## Acknowledgements

We are grateful to all individuals that participated in the research studies. We appreciate helpful discussions from all members of the Davis and Y. Chien labs. We also thank all staff members from the HIMC (Yael Rosenberg-Hasson, Michael D. Leipold and Weiqi Wang) for data analysis, management and helpful discussions, HIMC Biobank (Rohit Gupta and Janine Bodea Sung) for sample processing and storage, Stanford-LPCH Vaccine Program (Alison Holzer and Savita Kamble) for management of clinical studies. The Clinical and Translational Research Unit at Stanford University was supported by an NIH/NCRR CTSA award UL1 RR025744. This work was supported by NIH grants (U19 AI090019, U19 AI057229) and the Howard Hughes Medical Institute to M.M.D, and by the EU’s Horizon 2020 research and innovation program under the Marie Sklodowska-Curie grant (FluPRINT, Project No 796636) to A.T.

## Author contributions

A.T. downloaded data, coordinated the integration of the data into the FluPRINT database, advised on the database design and wrote the manuscript. I.T. built the MySQL database and wrote PHP script for the data import into the database, and contributed to writing the manuscript. H.T.M. managed the analysis, data collection and management of the SDM at the HIMC and advised during the manuscript preparation. C.L.D. was responsible for regulatory approvals, protocol design, study conduct, clinical data management and provided assistance during the manuscript preparation. M.M.D. conceived and supervised clinical studies and advised during the manuscript preparation.

